# Allantoin induces pruritus by activating MrgprD in chronic kidney disease

**DOI:** 10.1101/2020.10.26.354654

**Authors:** Yan Yang, Yulin Sun, Donglang Guan, Dan Chen, Dijun Wang, Tongtong Liu, Meixiao Sheng, Tao Jing, Shi Jun, Chan Zhu, Guang Yu, Xinzhong Dong, Zongxiang Tang

## Abstract

Chronic kidney disease is a disease with decreased, irreversible renal function. Pruritus is the most common skin symptom in patients with chronic kidney disease, especially in end-stage renal disease (AKA chronic kidney disease-associated pruritus [CKD-aP]); however, the underlying molecular and neural mechanism of the CKD-aP in patients remains obscure. Our data show that the level of allantoin increases in the serum of CKD-aP and CKD model mice. Allantoin could induce scratching behavior in mice and active DRG neurons; the calcium influx and the action potential were significantly reduced in DRG neurons of MrgprD KO or TRPV1 KO mice. U73122, an antagonist of PLC, could also block calcium influx in DRG neurons induced by allantoin. Thus, our results concluded that allantoin plays an important role in CKD-aP, mediated by MrgprD and TrpV1, in CKD patients.

## Introduction

Itch is a symptom of self-protection in the evolution of organisms and widely exists in human, non-human primates, cattle, dog, rodents and even fish [1–5]. It is defined an unpleasant sensation that provokes the desire to scratch, and is classified as acute and chronic [6, 7]. Chronic itch is manifested in many diseases of dermatological, systemic, neurological or psychogenic origin [8]. Systemic diseases with clinically significant pruritus include cholestatic liver disease, endocrine/metabolic disease, hematologic/lymphoproliferative disease, and chronic kidney disease [9–14].

Chronic kidney disease (CKD) or end-stage renal disease (ESRD) has become increasingly drastic over the past decades worldwide and has become a serious public health issue [15]. In patients with CKD, pruritus is a common and painful symptom. The clinical characteristics of CKD-associated pruritus patients vary greatly from patient to patient [13, 15]. Although pruritus in some patients is intermittent, pruritus in others is persistent and at night more severe than during the day [16, 17]. Pruritus seriously affects patients’ quality of life by sleep disturbance, anxiety, depression, skin injury, and infection [18–20]. A series of clinical methods have been adopted for the treatment of CKD-associated pruritus, such as rehydration emollients for topical treatment (tacrolimus ointments and gamma linolenic acid ointment) [21, 22], system treatments (gabapentin and pregabalin, naltrexone (a μ-opioid receptor antagonist), nalfurafine (a κ-opioid receptor agonist), pentoxifyline, thalidomide) [23–27], phototherapy with type B ultraviolet light (UVB), and acupuncture [28–30]. However, most patients with pruritus-related CKD are unresponsive to these treatments. This is mainly because the pathogenesis of CKD patients is clearly multifactorial and still poorly understood.

Itching substances produced by metabolic disorders caused by chronic kidney injury activate itch-related receptors, which cause pruritus in chronic kidney disease. Metabonomics studies have indicated significant differences between some substances in the blood of pruritus patients with CKD-related pruritus and normal plasma. Yasutoshi Akiyama et al. analyzed over 500 compounds, quantified 146 compounds, and identified that the plasma concentrations of 25 compounds were significantly changed, and suggested that four compounds (4-oxopentanoate, hippurate, allantoin and cytosine) may be common uremic compounds [31]. Among these compounds, increased levels of allantoin have been found in some diseases, including CKD, diabetic nephropathy, polycystic kidney disease (PKD), inflammatory autoimmune condition and kidney transplants [32–35]. Allantoin may promoted fibroblast proliferation and synthesis of the extracellular matrix [36]and is often used in various cosmetic lotions and creams, and pharmaceutical products [37]; however, some people who used these products reported itching symptoms. Therefore, we speculate that allantoin, as an intermediate in uric acid metabolism, may be a substance causing pruritus in chronic kidney disease. Our recent results showed that allantoin isolated from Yam could directly induce the scratch behavior and activated DRG neurons in mice [38].

Itch receptors can be classified into histamine-dependent and non-histamine-dependent receptors. However, antihistamine therapy in patients with pruritus of CKD is ineffective. Mas-related G protein-coupled receptors (Mrgpr) are composed of more than 50 members, mainly expressed in the peripheral nervous system and related to sensory functions [39]. So far, the main function of Mrgprs is itch-related function and non-histamine-dependent pruritus [9, 10, 40–42]. Hence, we hypothesize that Mrgprs are probably the receptors of metabolites in the metabolic process of chronic kidney disease. It is likely that allantoin may directly activate the pruritus receptors of Mrgprs to induce pruritus in chronic kidney disease. Based on this hypothesis, we tested the serum components of CKD in humans and chronic kidney injury in mice. As expected, allantoin in the serum of CKD and the kidney injury model in mice increased significantly, and could directly activate MrgprD in the Mrgprs family to induce itching behavior.

## RESULTS

### 1. Serum from pruritus patients with chronic kidney disease could induce itch behavior in mice

Pruritus is an important clinical symptom in patients with chronic kidney disease. To determine the substance that causes pruritus in chronic kidney disease, we used serums from patients and healthy people in behavioral experiments in mice, respectively. First, we used ultrafiltration centrifugation to filter macromolecular substances (>3 kd) in serum of the CKD patients with CKD and of healthy people. Then, we injected filtered serums from different groups into the necks of the mice. The results showed that scratching numbers induced by the serum of the patients with CKD were significantly more than in healthy people (52.25±2.81 vs.15.25±1.7, *P*<0.01, n=4) (**Figure 1A**). This suggested that there is really something in the patients’ sera that can induce itching in mice, and they are small molecules less than 3 kd. These small molecule substances are not only the cause of pruritus in patients with CKD, but also induce pruritus in mice.

**Figure 1.**
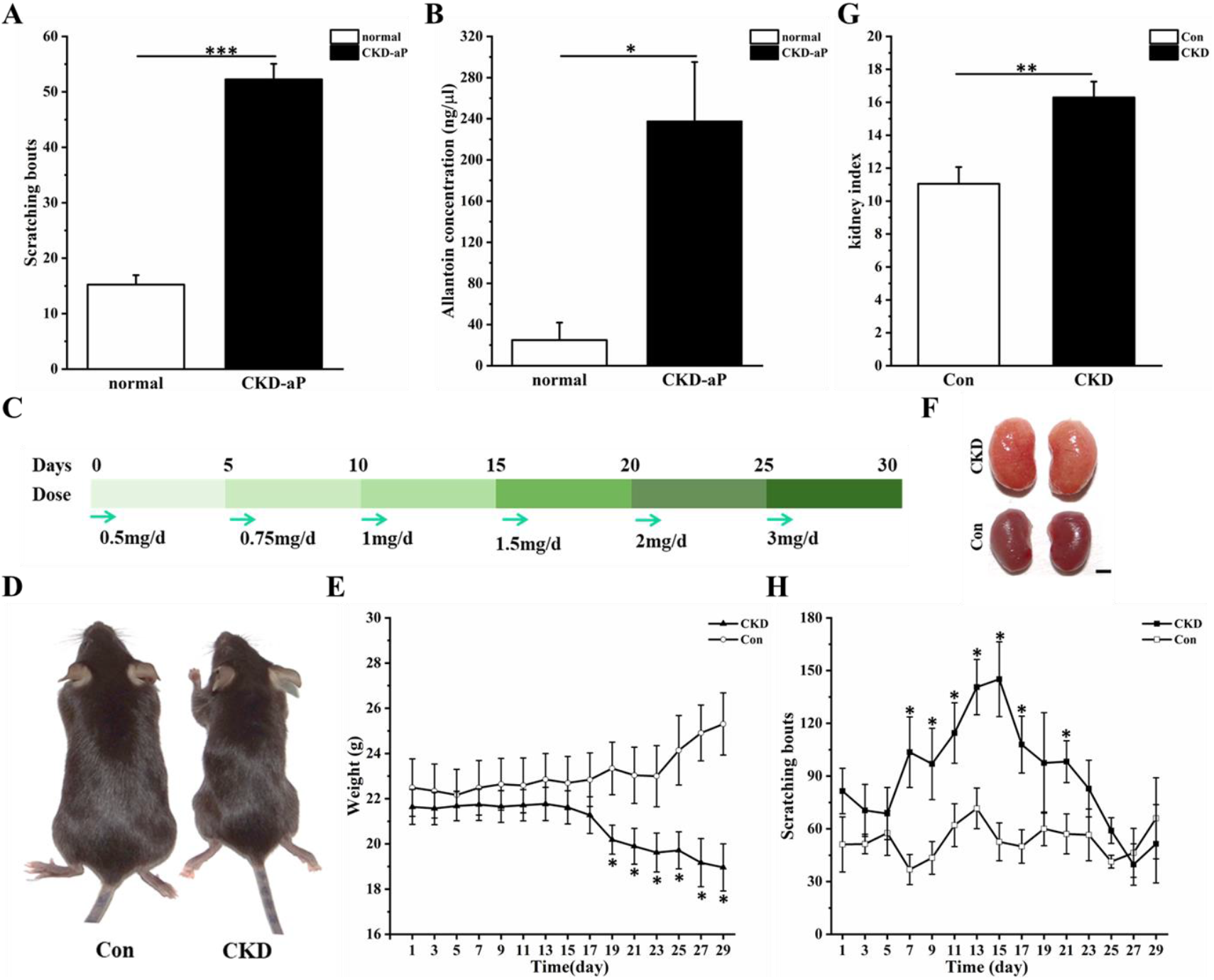
Scratching numbers of CKD model mice increased. **(A)** Small molecular (Mr<3Kd) in serum of the CKD-aP patients induced more scratching numbers compared with normal people(*p*<0.001). Data were expressed as mean ± s.e.m. **(B)** Allantoin concentration was higher in the serum of CKD-aP patients than normal people (*p*=0.022). **(C)** Scheme of treatment in adenine-induced mouse CKD. **(D-E)** Volume and weight of CKD model mice were significantly reduced than mice in control group (18.96 ± 1.04 vs. 25.30 ± 1.36, *p*<0.01, n=3, day29). **(F)** CKD model mice exhibited apparently renal injury compared with control mice (saline was given to the control mice). CKD model mice showed significant kidney edema. **(G)** Kidney index of the CKD model mice was conspicuously higher than that of control group (kidney index = kidney weight/weight). **(H)** Scratching numbers of the CKD model mice increased more than control group (the scratching bunts contained the neck and ears of the mice). All data are expressed as mean ± s.e.m. The data were statistically analyzed with two-tailed, paired/unpaired Student’s t test and a one-way or two-way ANOVA.

### 2. Allantoin in serum of patients with chronic kidney disease enhanced significantly

Patients with chronic nephropathy often have disorders of purine metabolism due to kidney injury. However, allantoin is a product of purine metabolism; the disorder of purine metabolism may lead to abnormal changes of allantoin content in the body. To detect the changes of allantoin in patients with CKD, the serum of chronic kidney patients and healthy people were collected from Jiangsu Provincial Hospital of Traditional Chinese Medicine. We performed UHPLC analysis using an ACQUITY UHPLC system (Waters Corporation, Milford, USA). The results showed that the allantoin concentration obviously increased in patients with CKD compared with the healthy people group (238±57 μg/ml vs. 18±13.5 μg/ml,*p* < 0.05, n=8) (**Figure 1B**), but other purine metabolites showed no significant changes (**Table 1**). Consistent with previous studies, the allantoin and urea in the serum of patients with CKD increased significantly rather than other purine metabolites [43, 44]. Therefore, allantoin may be a candidate to induce pruritus in nephropathy.

**Table 1.** Changes of purine metabolites in serum of chronic kidney disease patients and CKD model mice

|  |  |  |  |  |  |  |
| --- | --- | --- | --- | --- | --- | --- |
| Samples | CKD-aP patients(μg/ml) |  |  | CKD model mice(μg/ml) |  |  |
| Variable | Con | CKD | P value | Con | CKD | P value |
| adenine | 0.005 ± 0.002 | 0.014 ± 0.009 | 0.536 | 0.014 ± 0.006 | 0.025 ± 0.009 | 0.325 |
| xanthine | 0.168 ± 0.122 | 0.046 ± 0.023 | 0.175 | 1.885 ± 0.558 | 1.953 ± 0.407 | 0.921 |
| hypoxanthine | 0.661 ± 0.077 | 1.707 ± 0.417 | 0.132 | 0.026 ± 0.011 | 0.046 ± 0.015 | 0.325 |
| uric acid | 0.077 ± 0.050 | 0.049 ± 0.038 | 0.669 | 0.068 ± 0.018 | 0.066 ± 0.008 | 0.901 |
| allantoin | 0.018 ± 0.0135 | 0.238 ± 0.057 | *0.021 | 7.874 ± 0.893 | 22.683 ± 4.118 | **0.005 |
| allantoic acid | 0.146 ± 0.030 | 0.208 ± 0.098 | 0.654 | 0.535 ± 0.130 | 0.411 ± 0.098 | 0.458 |
| urea | 66.542 ± 8.108 | 136.226 ± 16.240 | *0.019 | 112.494 ± 9.215 | 216.995 ± 29.099 | **0.005 |

Data are expressed as mean ± SD for normally distributed continuous variables. *P* values are calculated using the 2-sample *t* test. The level of allantoin and urea increased in serum of CKD-aP patients and CKD model mice, compared with control group. (\**p*<0.05, \*\**p*<0.001).

### 3. The mouse model used in the study of chronic kidney disease showed obvious scratching behavior

Many experimental animal models have been used to study CKD. The traditional CKD model uses Sprague-Dawley (SD) rats treated with adenine intragastric and administers 200mg/kg/d or five-sixths nephrectomy. However, these methods had a very high mortality rate in rats or were relatively resistant to hypertension and decline in renal function in C57BL/6 mice [45]. To imitate the CKD in C57BL/6 mice, similar to the method of the rat model establishment, mice were also treated with adenine intragastric administration 200mg/kg/d; the same outcome occurred: the normal survival rate was only 33.3%(n=6). To reduce the mortality rate of the model mice, changes have been made to the method of model building. Adenine concentration, which is mainly induced by nephropathy, was gradually increased. The initial concentration of adenine was 0.5mg/ml with an adding gradient every five days (**Figure 1C)**. After the model was established, the weight of the model animals was reduced significantly more than the index of the control group at the 19^th^ day (20.19±0.64 vs. 23.34±0.64, *p*<0.05, n=10) (**Figure 1D, E**). The renal index of the model group increased significantly compared with the control group (16.3±0.96 vs. 11.0±1.0, *p*<0.01, n=10) (**Figure 1F, G**). The scratching behavior of animals was also observed and analyzed in the process of model building. The scratch numbers of the mice in model group at the 7^th^ day were more than the numbers in control group (111.86±26.49 vs. 36.87±8.56,*p*=0.013, n=10) (**Figure 1H)**. Following the development of the CKD model mice, the number of scratches increased gradually and reached its maximum on the 15^th^ day. With the continuation of the kidney injury model, the activity, weight, and scratch numbers of the model mice decreased obviously. Finally, the mice died of severe kidney damage.

To observe the kidney damage caused by the high purine feeding, kidney tissues were obtained at different stages of mouse modeling on day 5, day 15 and day 30, respectively. Renal fibrosis and inflammatory cells were analyzed by hematoxylin-eosin staining. The results indicated that the fibrosis area was markedly increased in the CKD model group on day 15 and day 30 compared with the control group (8.61±0.46%, day-15; 27.82±2.91%, day-30; vs. 0.37±0.21%, *p*<0.01, n=3), and that the inflammatory cell infiltration was also significantly enhanced in the CKD group mice compared with control mice (144±6 inflammatory cells/HPF, day 15; 565±38 inflammatory cells/HPF, day 30; vs. 51±10 inflammatory cells/HPF, *p* <0.01, n=3) (**Figure 2A, B, C**).

**Figure2.**
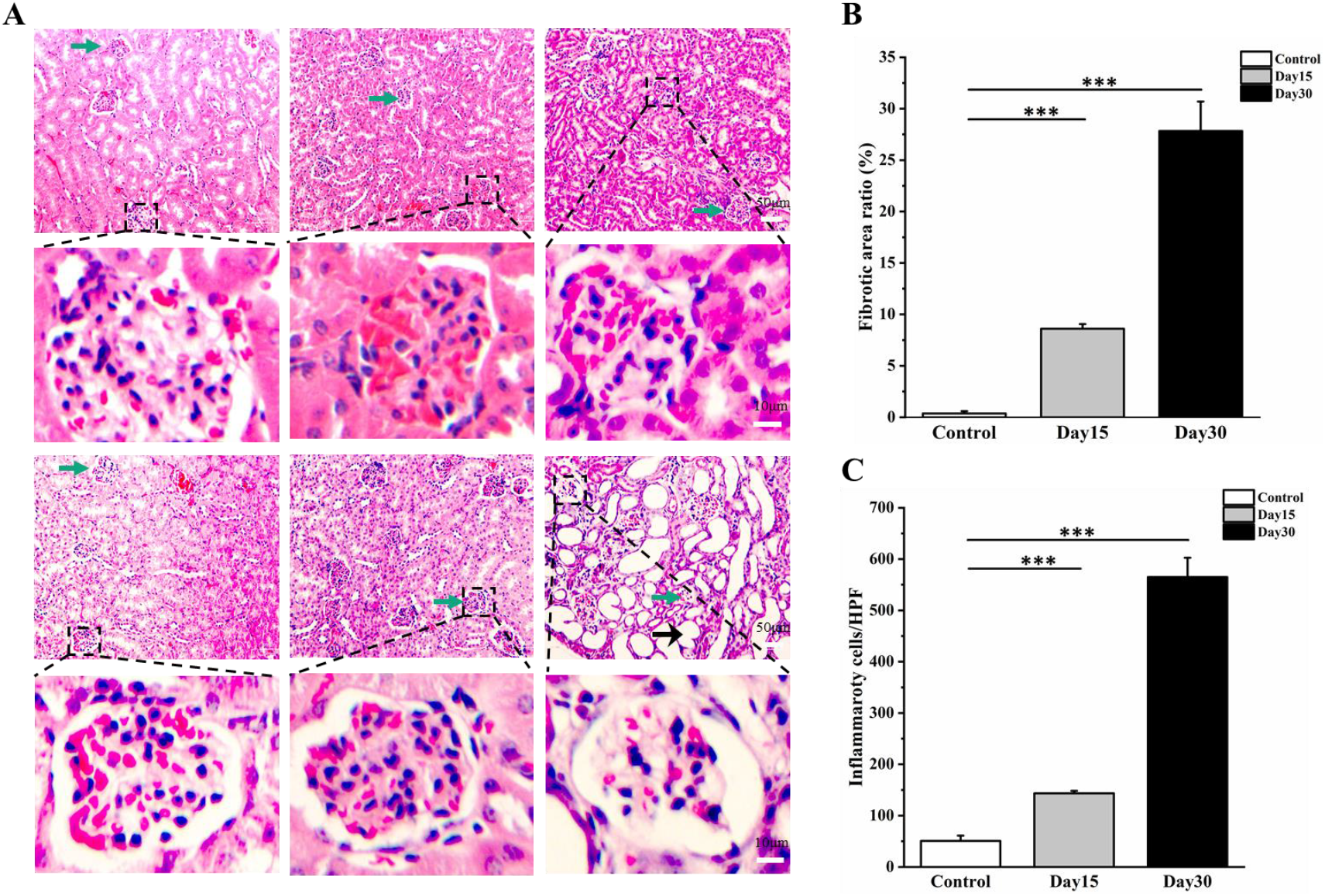
CKD model showed obvious renal injury. **(A)** Representative images of kidney stained by in CKD model mice showing distinct renal injury compared with the control group, day15, day30 respectively. Green arrow indicated the glomerulus. Black arrow indicated the renal fibrosis. Dotted boxes represented the locally enlarged areas. Scale bar, 50μm or 10μm. **(B)** CKD model mice exhibited serious renal fibrosis in 15^th^-day and 30^th^-day compared with the control mice (\*\*\**p*<0.001). **(C)** Inflammatory cell infiltration also was significantly enhanced in CKD group mice in 15^th^-day and 30^th^-day compared with control mice (\*\*\**p*<0.001).

### 4. Allantoin levels of serum, kidney and urine samples in model mice increased significantly

Allantoin in serum samples of patients with CKD has been determined to be significantly higher than that of healthy people controls. To confirm that allantoin levels in the kidney injury mice model were similar to those in patients, tissues and samples (serum, urine and kidney tissue) from model mice were collected and the components of these samples were analyzed. The allantoin and urea in the serum of model mice were significantly higher than those of control group (22.68±4.12 vs. 7.87±0.89, n=8, *p*<0.01 and 216.99±29.1 vs. 112.49±9.22, n=8, *p*<0.01). Adenine and hypoxanthine in kidney tissue decreased (2.822±0.46 vs. 1.73±0.27, n=8, *p*<0.05 and 6.76±1.82 vs.3.16±0.49, n=8, *p*<0.05); the allantoin increased significantly (16.22±1.96 vs. 49.16±10.09, n=8, *p*<0.05). Although adenine decreased in urine samples, no changes were detected in allantoin and other metabolites (**Table 2**). The concentration of allantoin in serum increased in patients with CKD and in the injured model mice. This means that kidney injury causes the allantoin metabolism pathway disorder, and then causes various abnormal physiological reactions.

**Table 2.** Changes of purine metabolites in kidney and urine of CKD model mice

|  |  |  |  |  |  |  |
| --- | --- | --- | --- | --- | --- | --- |
| Samples | Kidney(μg/ml) |  |  | Urine(μg/ml) |  |  |
| Variable | Con | CKD | P value | Con | CKD | P value |
| adenine | 2.822 ± 0.460 | 1.726 ± 0.270 | *0.035 | 0.642 ± 0.133 | 0.331 ± 0.078 | *0.04 |
| xanthine | 17.647 ± 1.389 | 19.109 ± 1.383 | 0.473 | 0.929 ± 0.230 | 0.618 ± 0.125 | 0.208 |
| hypoxanthine | 6.763 ± 1.821 | 3.155 ± 0.493 | *0.034 | 0.946 ± 0.300 | 0.506 ± 0.150 | 0.157 |
| uric acid | 0.789 ± 0.121 | 0.603 ± 0.052 | 0.124 | 0.286 ± 0.057 | 0.274 ± 0.024 | 0.822 |
| allantoin | 16.219 ± 1.955 | 49.1556 ± 10.089 | *0.01 | 209.216 ± 38.125 | 214.750 ± 25.331 | 0.901 |
| allantoic acid | 4.765 ± 0.584 | 5.299 ± 0.555 | 0.532 | 10.086 ± 6.845 | 15.834 ± 3.419 | 0.407 |
| urea | 206.701 ± 22.224 | 273.293 ± 31.833 | 0.124 | 793.800 ± 129.776 | 699.345 ± 61.400 | 0.459 |

Data are expressed as mean ± SD for normally distributed continuous variables. *P* values are calculated using the 2-sample *t* test. The level of allantoin and urea increased in serum of CKD-aP patients and CKD model mice, compared with control group. The level of allantoin and urea increased in serum of CKD model mice, comparing with control group (\**p*<0.05, \*\**p*<0.001).

### 5. Allantoin could induce scratching behavior in mice

The serum of patients with CKD could induce pruritus in mice. Moreover, allantoin concentration in serum of patients with CKD was obviously higher than that of healthy people. It is quite possible that allantoin is the substance that induces inching. To prove that allantoin is an itchy substance in pruritus of CKD, allantoin (10mM, 50μl) was injected subcutaneously into the neck of the mice. As we assumed, allantoin could induce more scratching numbers than control group (62±5.68 vs. 26.6±8.51, *p*<0.01, n=5) (**Figure 3A**). However, scratching bouts induced by urea showed no difference compared with control mice (**Figure 3B**, 33±4.04145 vs. 41±1.73205, *p*=0.284). To further prove the itching effect of allantoin, allantoin was also injected into the tail vein of mice to increase allantoin concentration in the veins of mice; the mice showed more scratching behavior compared with normal mice (97±6.24 vs. 46±5.66, *p*<0.001, n=5) (**Figure 3C**). If other itching substances were injected into the tail vein, the scratching behavior induced by chloroquine was significantly higher than that of the control group (81±10.1 vs. 41±7.1, *p*<0.05, n=3) (**Figure 3D**), histamine also caused more scratching reaction (103±1.5 vs. 41±7, *p* <0.01, n=3) (**Figure 3D**). These results clearly indicate that allantoin, chloroquine and histamine tail vein injection could evoke scratching reaction in mice. Allantoin has the same itch-inducing function as chloroquine and histamine.

**Figure3.**
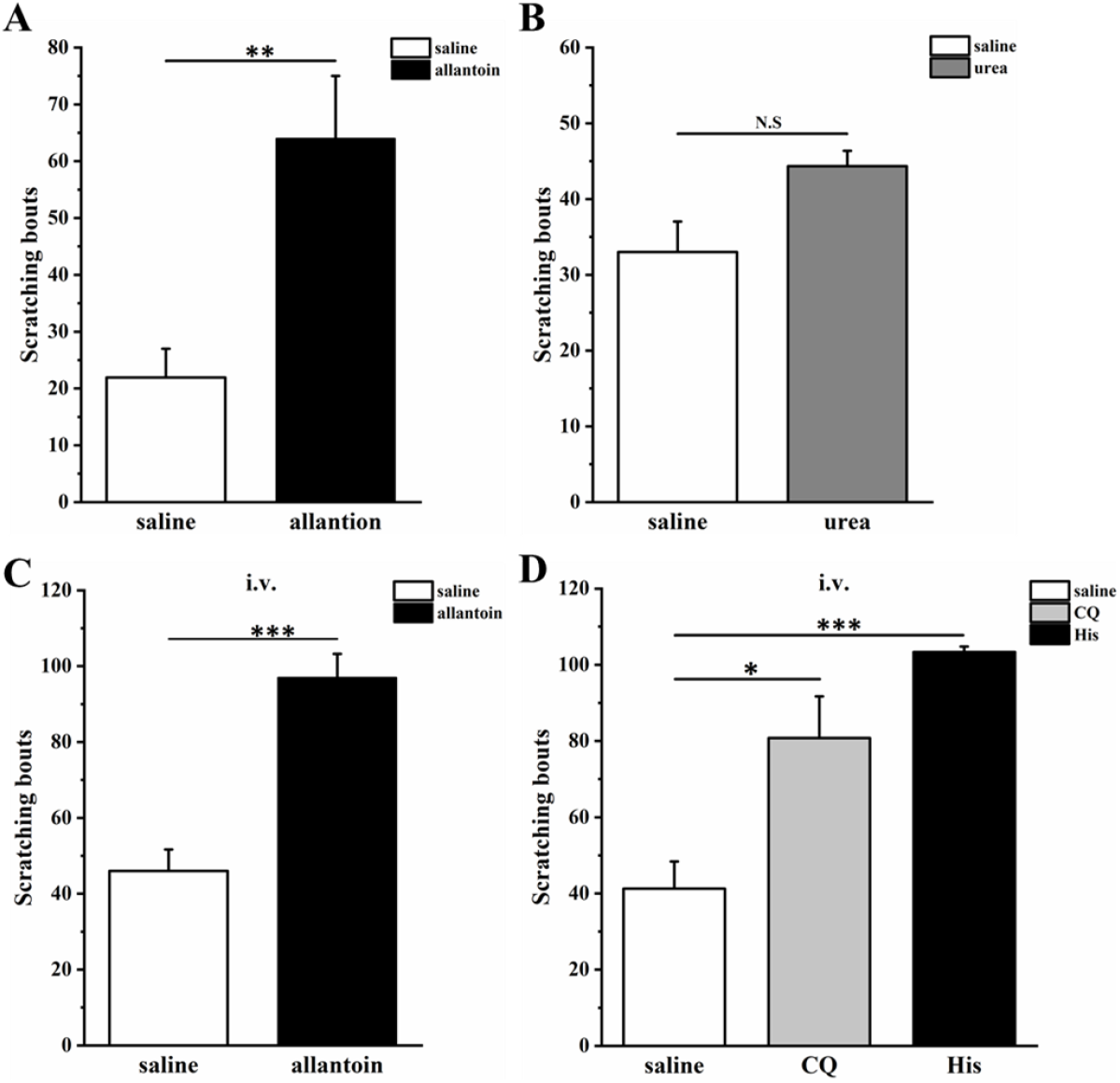
Allantoin induced pruritus behavior in mice. **(A)** Allantoin with subcutaneous injection in the neck of the mice induced more scratching bunts than saline group (\*\**p*=0.001). **(B)** Scratching numbers showed no different between urea group and control group. **(C)** Allantoin with intravenous also induced more scratching numbers compared with control group (*p*=0.28, n=3). **(D)** CQ and histamine with intravenous induced more scratching numbers than control group respectively. All data are expressed as mean ± s.e.m.

### 6. Allantoin could activate cultured DRG neurons

Since allantoin has been considered a candidate for itching induction, next we wanted to know whether it could activate neurons directly. We adopted calcium imaging and electrophysiological recording techniques to determine the function of allantoin. When allantoin was added to a chamber buffer, DRG neurons were evoked to produce calcium influx (**Figure 4A, B**). DRG neurons activated by allantoin were mainly small diameter neurons ranging in size from 11 to 23 microns (**Figure 4C**). These activated neurons accounted for about 9.1% of the total statistical neurons (454/5005) and were dose-dependent; the EC50 of allantoin was 0.5mM (**Figure 4D, E**). Then, we used a patch clamp technique to record these small diameter neurons; the results indicated that allantoin could directly activate these DRG neurons to evoke action potentials (**Figure 4F, G**). Therefore, we deduced that allantoin activated itch-related neurons and resulted in itching behaviors in humans and mice.

**Figure4.**
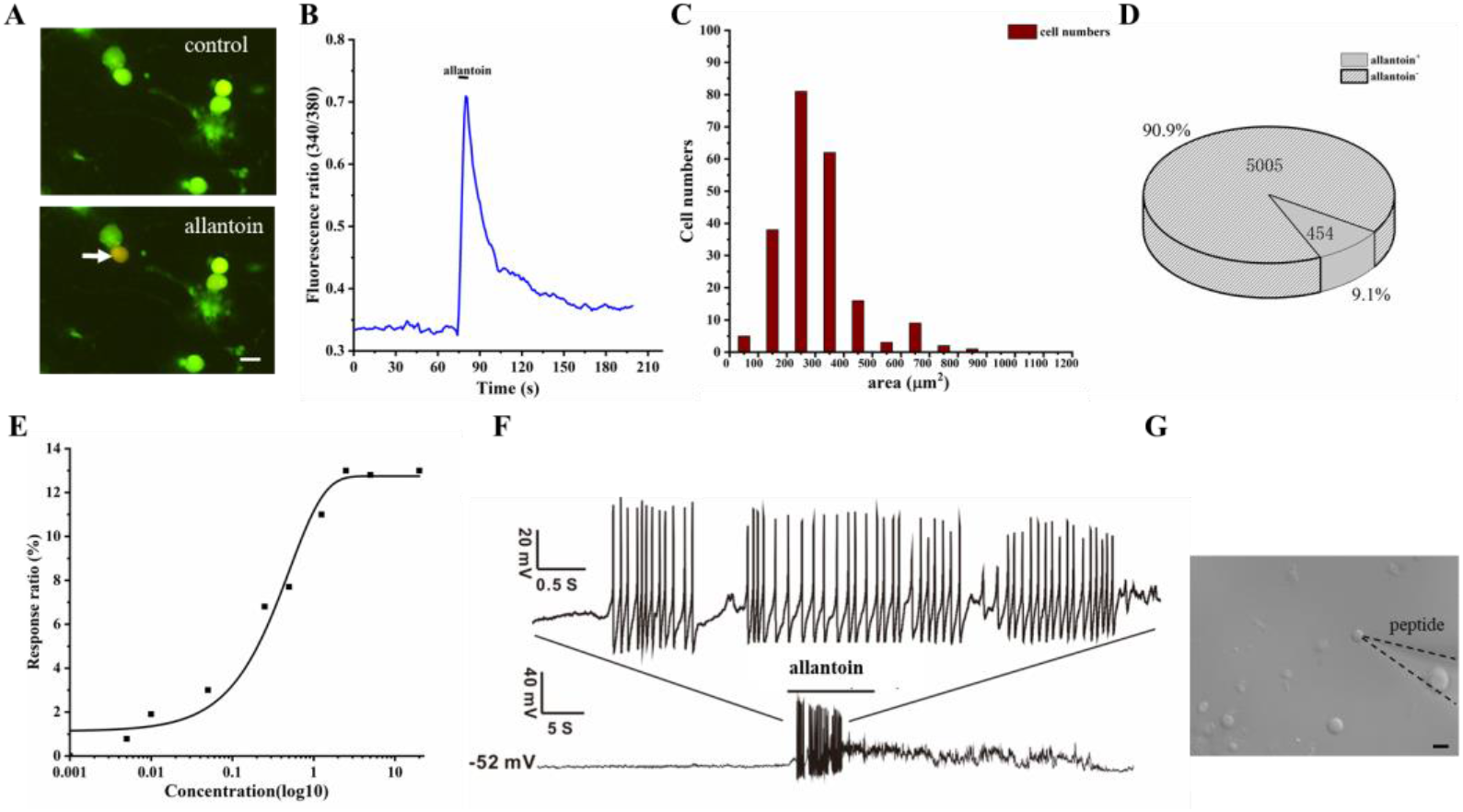
Allantoin induced calcium incurrent and action potential in DRG neurons. **(A)** Allantoin could activate the DRG neurons. The flat arrow indicated the allantoin positive DRG neurons. **(B)** Fluorescence ratio of the DRG neurons induced by allantoin. **(C)** Allantoin activated small and medium diameters of DRG neurons **(D)** 9.1% DRG neurons was could activated by allantoin in cultured neurons (allantoin activated positive neurons/total neurons, 454/5005). **(E)** EC50 of the allantoin was 0.5mM. **(F)** Photo showed the DRG neuron recorded AP Dotted line shows the location of the recording electrode. **(G)** Allantoin induced action potential in DRG neurons (n=6). Scale bar 20μm.

### 7. MrgprD is a target of allantoin-induced pruritus

In previous studies, MrgprA3, MrgprA1, MrgprC11, MrgprD and MrgprB2 in the Mrgprs family were involved in the formation of itching sensation [10, 40–42, 46–48]. Except for MrgprB2, other itch receptors of Mrgprs were only expressed on DRG neurons [39]. Allantoin could activate DRG neurons and induce itching behaviors; however, the report of clinical results showed that antihistamines were ineffective against pruritus in patients with CKD. Therefore, the pruritus receptors in the Mrgprs family were the most likely target candidates. To find the relationship between allantoin and Mrgprs receptors, a Mrgpr cluster KO (Mrgpr-clusterΔ^-/-^) and MrgprD KO mice were used for testing allantoin-activated receptors. When allantoin was injected into the Mrgpr cluster KO and WT mice, allantoin-induced pruritus showed almost no difference (86 ± 8.08 vs. 72.3 ± 11.8, *p*=0.393, n=3) (**Figure 5A**), but when allantoin was injected into the MrgprD KO and WT mice, allantoin-induced pruritus in MrgprD KO mice was significantly lower than in control group (WT) (56.9 ± 5.2 vs. 26.7 ± 5.8, *p*<0.001, n=10) (**Figure 5B**). Next, we examined the responses of allantoin to DRG neurons; allantoin-sensitive DRG neurons could also be activated by β-alanine (**Figure 5C, D**). These activated neurons were all MrgprD-positive neurons, but neurons without GFP markers (MrgprD negative) almost did not respond to allantoin (**Figure 5E**). The response intensity of DRG neurons induced by allantoin and β-alanine was almost the same (**Figure 5E)**. About 83% β-alanine-sensitive neurons could also be activated by allantoin (**Figure 5G**). To further verify that allantoin could activate the MrgprD receptors, alamandine of another MrgprD agonist was applied to the cultured DRG neurons [49, 50]. About 88% allantoin-sensitive neurons could also be activated by alamandine and β-alanine **(Figure 5F, H, I);** however, the response of DRG neurons to allantoin in MrgprD KO mice was deficient (**Figure 5J, K**). These results suggest that MrgprD is probably a target for allantoin and played an important role in itch sensation induced by allantoin.

**Figure5.**
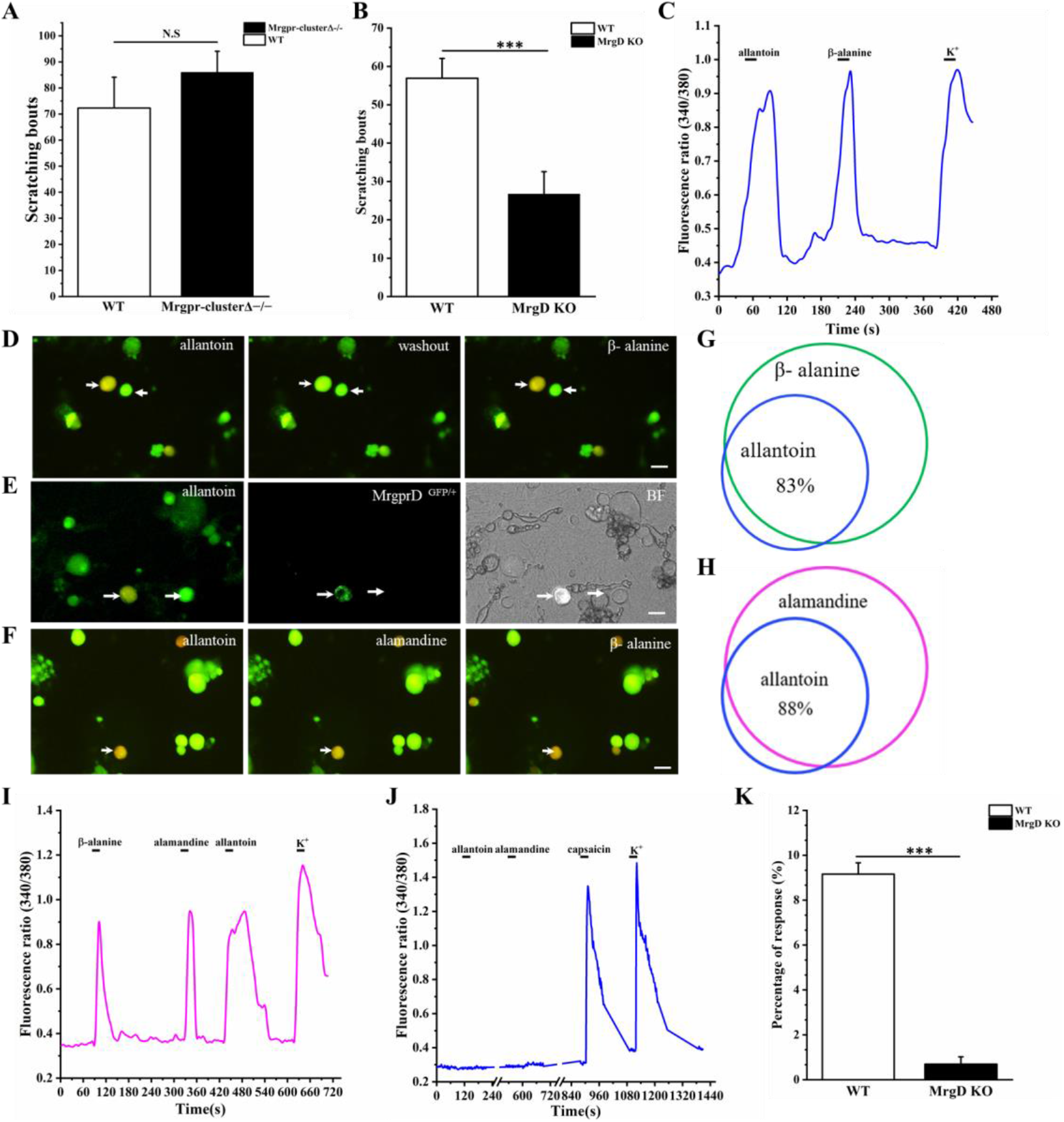
MrgprD participates in allantoin-induced scratching behavior in mice and the agonists of MrgprD activate the DRG neurons activated by allantoin. **(A)** Scratching numbers in Mrgpr Cluster KO mice was less than wide type mice (p=0.39). **(B)** MrgprD KO mice showed obviously different in scratching numbers compared with wide type mice (27 ± 6, n=10, *p*<0.001). **(C)** Fluorescence ratio of DRG neurons activated by allantonin. **(D)** Allantoin (0.5mM) positive neurons could activated by β-alanine (0.5mM). Swallowtail arrow indicates allantoin positive neuron, flat-ended arrow indicated negative neuron. **(E)** Allantoin positive neurons was also MrgprD positive neuros (MrgprD^GFP/+^ neurons). Swallowtail arrow indicates MrgprD positive neuron, flat-ended arrow indicated negative neuron. BF means bright field. Scale bar, 20μm. **(F)** The same DRG neuron could activated by allantoin, β-alanine and alamandine (10μM) (MrgprD antagonist). Scale bar, 20μm. **(G)** Allantoin positive DRG neurons showing obvious calcium ion influx could activated by β-alanine. Blue circle indicated allantoin positive neurons, and green circle indicated β-alanine positive neurons. **(H)** Allantoin positive DRG neurons showing obvious calcium ion influx could activated by alamanine. Blue circle indicated allantoin positive neurons, and pink circle indicated β-alanine positive neurons **(I)** DRG neuron indicated by swallowtail arrow in (a) activated by both allantoin, β-alanine, alamanine **(J)** Allantoin and alamandine could not induced calcium ion influx in DRG neurons of MrgprD KO mice, while capsaicin still can activate DRG neurons. **(K)** Compared with the wild type mice, the percentage of positive DRG neurons induced by allantoin reduced in MrgprD KO mice.

### 8. TRPV1 is involved in allantoin-induced itch

TRPV1 and TRPA1 are two important ion channels in the TRPs family associated with pruritus, including histamine-dependent and non-histamine-dependent itching. To confirm whether TRPV1 and TRPA1 are involved in allantoin-induced itch, TRPV1 KO mice and TRPA1 KO mice were used to detect the itch-inducing function of allantoin. The results showed that scratching bouts in TRPA1 KO mice had no significant difference compared with *WT* mice (56.0 ± 10.5 vs. 51.8 ± 2.8, *p*=0.89, n=3) (**Figure 6A**), but TRPV1 KO mice exhibited significantly less scratching than *WT* groups (75.2 ± 15.0 vs. 27.6 ± 7.0 bouts, *p*<0.05, n=5) (**Figure 6B**). Next, DRGs from *WT* mice were collected, digested and cultured; then, 100μM capsaicin and 0.5mM allantoin were applied to these cultured DRG neurons. The neurons responding to capsaicin also responded to allantoin, and their co-reaction ratio reached 92% (**Figure 6C-E**). If in TRPV1 KO mice, the response of capsaicin- and allantoin-induced DRG neurons disappeared (**Figure F-G**). Ruthenium Red (10 μM), a TRP channels antagonist, could reduce the fluorescence ratio of DRG neurons induced by allantoin (0.53077±0.06542 vs. 0.06154±0.02412, *p*<0.001) **(Figure 6H).** Together, these data indicate that TRPV1 is required for allantoin-induced scratching in mice.

**Figure 6.**
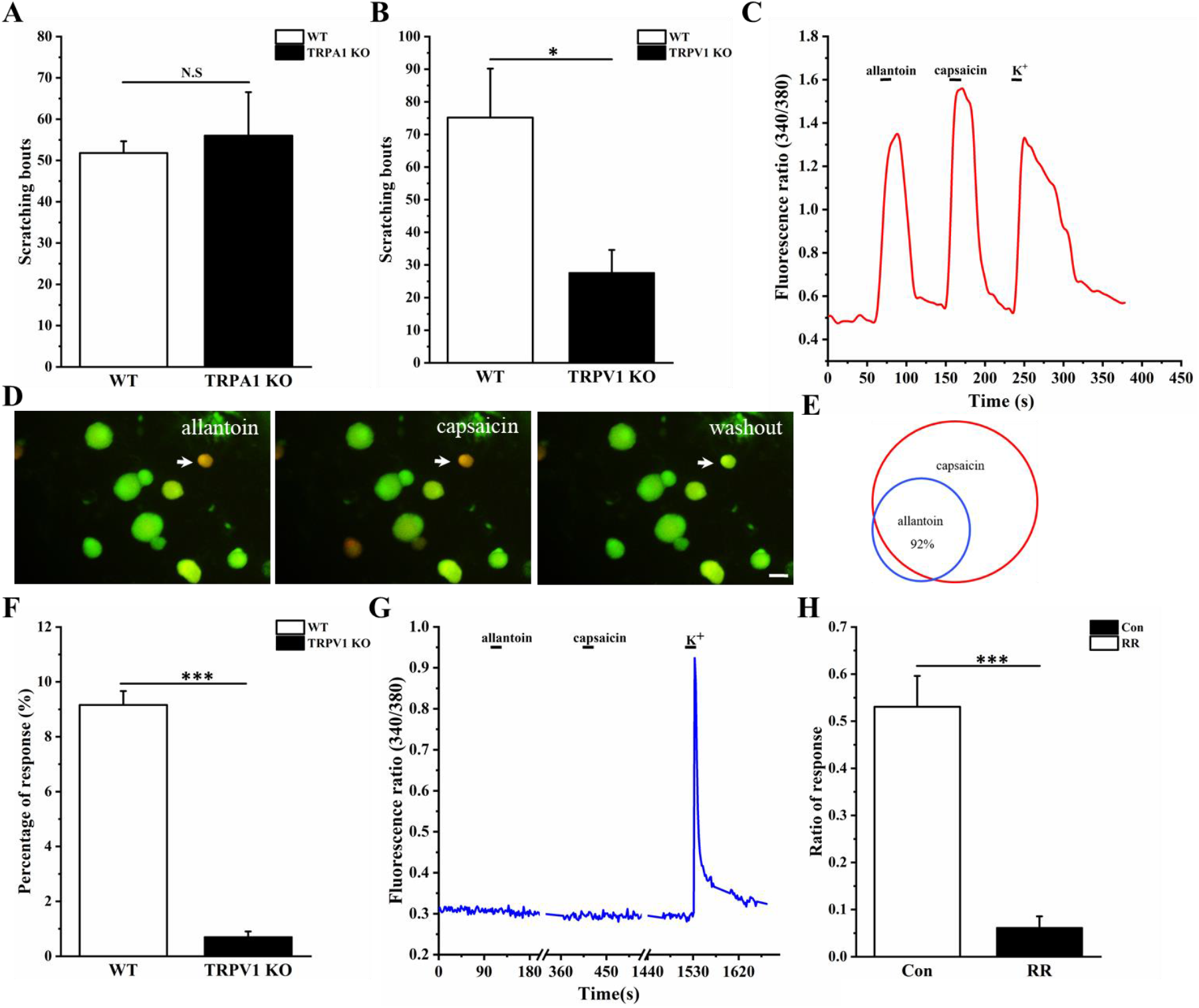
TRPV1 mediated scratching behavior induced by allantoin. **(A)** TRPA1 KO mice showed no obvious difference compared with control group (*p*=0.89). **(B)** Scratching numbers of TRPV1 KO mice reduced significantly compared with wild type mice (*p*<0.05). **(C)** Allantoin activated the TRPV1 positive DRG neuron. **(D)** DRG neuron indicated by the arrow can be activated by both allantoin and capsaicin to induce calcium influx. **(E)** 92% DRG neurons activated by allantoin could activated by capsaicin. Scale bar, 20 μm. **(F)** Allantoin could not induce calcium influx in DRG neurons of TRPV1 KO mice. The percentage of the positive neurons reduced obviously compared with wild type mice. **(G)** Responsive curve of DRG neurons of TRPV1 KO mice. **(H)** Ruthenium red (RR), an antagonist of TRPs channel, could reduce the responsive ratio of DRG neurons induced by allantoin(*p*<0.001).

### 9. The function of MrgprD requires the participation of TRPV1

Capsaicin and allantoin could activate the same DRG neurons, and the proportion of co-reaction is more than 90%. This indicates that most of MrgprD and TRPV1 may co-express on the same neurons [51]. To prove the functional relationship between MrgprD and TRPV1, relevant plasmids were co-transferred or transferred to HEK293T cells, respectively. Meanwhile, the blank plasmid was also transferred into the same cell line as a control. HEK293T cells transferred with blank plasmid did not respond to capsaicin and allantoin stimulation (**Figure 7A, B**). If the plasmid of MrgprD or TRPV1 was transferred separately; single MrgprD-transferred cells did not respond to allantoin stimulation, (**Figure 7A, C**), TRPV1-transferred cells only responded to capsaicin, but not to allantoin (**Figure 7A, D**). If the plasmids of MrgprD and TRPV1 were co-transferred, all of them (including allantoin, β-alanine and capsaicin) could induce the transferred cells to responds (**Figure 7A, E**). These data indicate that the functional relationship between MrgprD and TRPV1 does exist. It is likely that allantoin activates MrgprD and then induces TRPV1 excitation by downstream signaling pathways.

**Figure 7.**
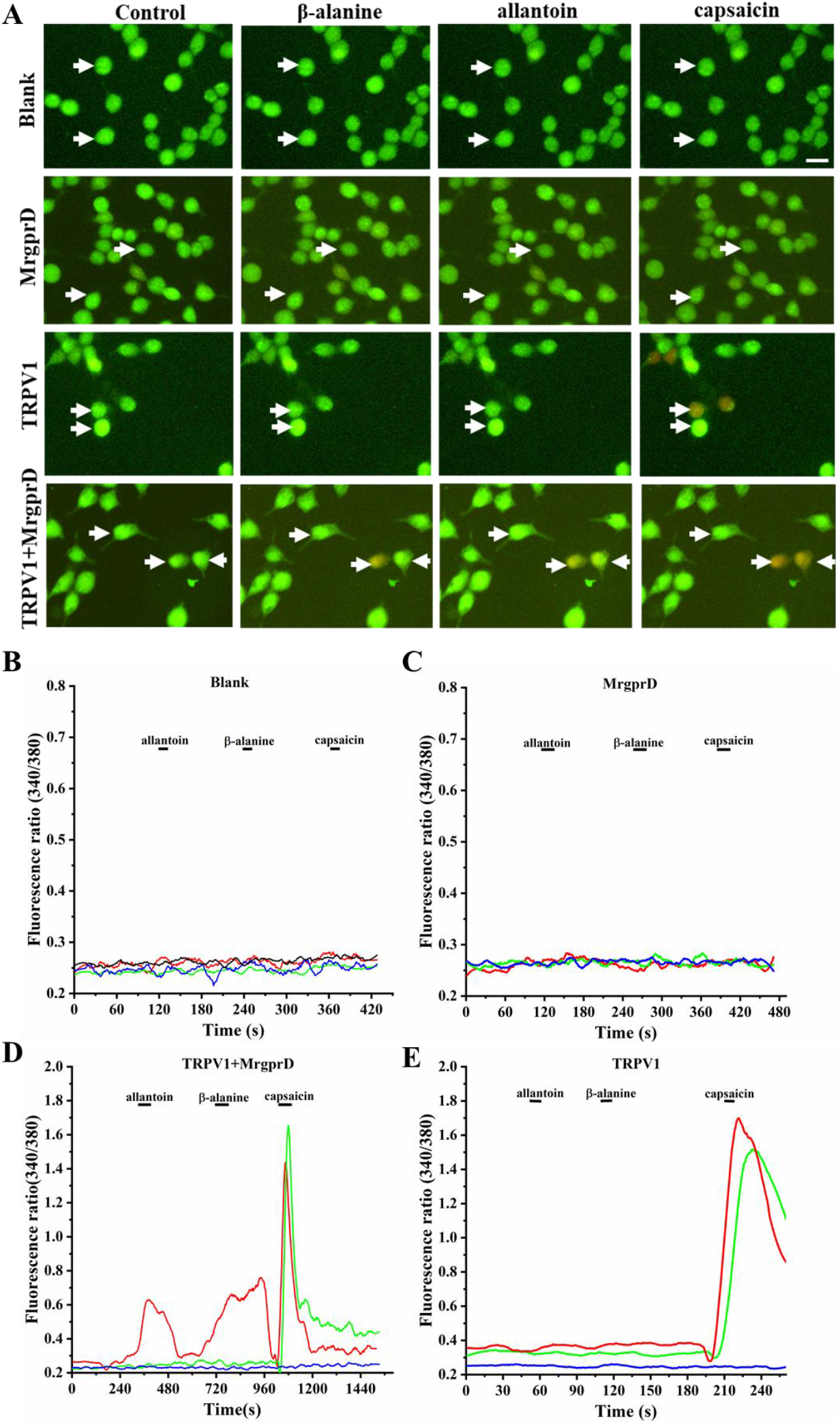
Allantoin activated HEK293T transfected with TRPV1 and MrgprD. **(A)** Representative images of β-alanine, allantoin-induced calcium imaging. HEK293T cells were transfected with MrgprD, TRPV1, or MrgprD companied with TRPV1. Allantoin, β-alanine induced calcium influx only in HEK293T cells transfected with MrgprD companied with TRPV1 compared with blank group (same treatment without plasmid). Capsaicin induced calcium influx only in TRPV1 and MrgprD companied with TRPV1. The swallowtail arrow indicated the positive cells only activated by capsaicin. The flat arrow indicated the positive cells activated by allantoin and capsaicin. **(B-C)** HEK293T cells showed no response. **(D-E)** The responsive curve of HEK293T cells induced by allantoin and β-alanine.

### 10. The functional coupling of MrgprD and TRPV1 is realized through the PLC pathway

As shown above, allantoin could activate DRG neurons and induce scratching behavior in mice, but if excluding MrgprD or TRPV1, the scratching behavior of mice and DRG neurons excitation disappeared. Indeed, there are some functional connections between MrgprD and TRPV1; however, the signaling pathways between MrgprD and TRPV1 remained elusive after allantoin was used as an agonist to activate MrgprD. To find out how the activation of MrgprD affects the function of TRPV1, we tested the effects of an inhibitor to the PLC pathway on the allantoin-induced calcium influx.

When cultured DRG neurons were pretreated with U73122 (a PLC selective antagonist), the allantoin-induced calcium influx was significantly less than that of the control group (0.51±0.08 vs. 0.27±0.07, *p*<0.001, n=13) (Fig 8.A). The reaction traces showed that the allantoin-induced response was obviously inhibited by U73122, then recovered after U73122 was removed (Fig 8. B). The above results indicate that the PLC inhibitor plays a major role in the allantoin-activated downstream signaling pathway of MrgprD in DRG neurons, and were indeed consistent with those reported in other studies [52] and that MrgprD by the PLC pathway induced calcium influx activated by β-alanine in Xenopus oocytes [53].

**Figure 8.**
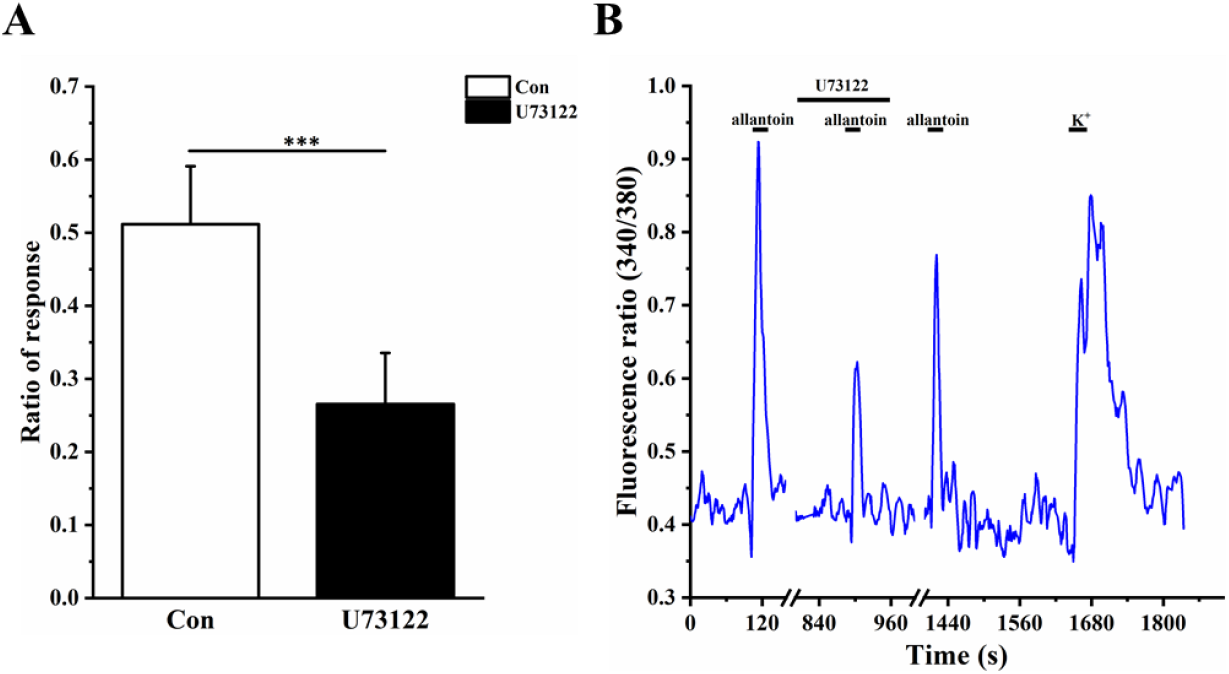
PLC signal pathway participates in allantoin induced calcium influx in DRG neurons. (A) Compared with the control group, PLC blocker U73122 (10μM) decreased the intensity of response induced by allantoin in DRG neurons. (B) U73122 can effectively inhibit allantoin-induced calcium influx in DRG neurons. *\*\*\*p<0.001.*

## Discussion

### 1. Allantoin was identified as an itchy substance in pruritus of chronic kidney disease

Pruritus is a common clinical symptom in patients with CKD; however, in earlier studies, the substances causing itching were unclear. In this study, we confirmed that allantoin not only increased significantly in the serum of patients with CKD, but also in the serum and kidney tissue of mice with chronic renal failure. These results are consistent with earlier reports [54, 55]. Importantly, our study found that allantoin, which changed in CKD, is actually an itch-causing substance. Because allantoin is an important intermediate product in purine metabolism, pruritus symptoms may occur in diseases caused by the disorder of purine metabolism. Therefore, it can be inferred that the pruritus caused by the disorder of purine metabolism may be caused by the change of the content of allantoin in vivo. This may not only explain the causes of pruritus in CKD, but also provide valuable clues for the study of the pruritus mechanisms of other purine metabolic disorders.

### 2. MrgprD is a receptor for pruritus in chronic kidney disease

In the clinical treatment of pruritus in CKD, there is almost no effective treatment, including antihistamines drugs. This brings great difficulties to clinicians in treating pruritus symptoms of chronic nephropathy, and also poses new challenges and research directions for researchers. Although MrgprD is an itch receptor that can be activated by β-alanine [42], this is the first time it has been found that MrgprD can be activated by allantoin and induce itching behavior. As a target of pruritus, MrgprD not only helps us understand the mechanism of pruritus formation in chronic nephropathy, but also provides a new way for the treatment of pruritus in chronic nephropathy. Moreover, this discovery also increases our understanding of the pruritus receptor MrgprD, which can be activated by allantoin in addition to β-alanine. This may partly explain why antihistamines are ineffective in the treatment of pruritus in CKD.

### 3. TRPV1 is the downstream ion channel of allantoin-activated MrgprD-induced itch

TRP channels are more diverse in activation mechanism and function selection than others in channels, and have been shown to play crucial roles in temperature, mechanosensation, olfaction, vision, taste and pain [56, 57]. More and more studies have shown that TRP channels are involved in the formation of itching sensation. TRPV1 is the downstream ion channel of histamine-induced itch, the PLCβ/PKC and PLA2/lipoxygenase pathways that have been considered to link histamine receptors to TRPV1 [58–61]. In histamine-independent itch, TRPA1 is the downstream ion channel of some of the stimulated itch receptors, such as MrgprA3, TGR5 and 5-HT7 that are Gβγ dependent while MrgprC11 and TSLPR signaling require PLC [62–65]. Moreover, in other histamine-independent itch receptors, the downstream signaling pathways require both TRPV1 and TRPA1, bradykinin causes itching by activating B2R and requiring downstream TRPV1 and TRPA1 [66, 67]. Leukotriene B4, an inflammation related leukotriene, requires both TRPV1 and TRPA1 downstream of LTB4 receptors 2 to generate itch [68]. Interleukin-31 (IL-31) is an endogenous factor in inflammatory and lymphoma-associated itch, an IL-31 receptor (IL-31RA) co-expressed with TRPV1 and TRPA1 on a small subpopulation of DRG neurons, and plays a crucial neuron-immune link role between TH2 cells and sensory neurons [69]. In our study, allantoin, a metabolite produced in CKD, was found to activate the histamine-independent pruritus receptor MrgprD, which then stimulated the downstream TRPV1 channel to produce itching sensation. This study not only clarified the causes of pruritus in chronic kidney disease, but also offers a direction for the treatment of pruritus symptoms in CKD. TRPV1 may become an important candidate receptor for the treatment of pruritus in CKD in the future.

In conclusion, allantoin is an important itchy substance that causes pruritus in CKD. Allantoin activated the pruritus receptor of MrgprD and became a new agonist of the receptor. These new findings are of great significance to the formation mechanism and clinical treatment of pruritus in chronic kidney disease. In addition, MrgprD activates the TRPV1 channel downstream through the PLC signaling pathway to participate in itching formation, which not only deepens our understanding of the relationship between G-protein-coupled receptors and ion channels, but also provides another way for the clinical treatment of pruritus in CKD. TRPV1 channel blockers may be an important choice in the future clinical treatment of pruritus in CKD.

## Materials and Methods

### Animals

Adult C57BL/6 mice weighing 18-20g were kept in a constant temperature and humidity environment with a standard 12-h light/dark cycle. All procedures for this study were approved by the Institutional Animal Care and Use Committee of the Nanjing University of Chinese Medicine.

### Chronic kidney disease models

Mice were randomly divided into two groups. Mice were given adenine at regular intervals twice a day. Control mice were given 0.9% saline. The dose of adenine increased gradiently every 5 days (0.5mg/ml, 0.75mg/ml, 1mg/ml, 1.5mg/ml, 2mg/ml, 3mg/ml).

### DRG neuron culture

Adult C57BL/6 mice were euthanized with isoflurane. DRGs were collected in DH10 with 10% Fetal Bovine Serum (Biological Industries, Israel), 90% DMEM/F-12(BI, Israel),1% penicillin-streptomycin (Solarbio, China) and then washed with HBSS. DRGs were dispersed in enzyme solution with 3.5 mg/ml dispase II, 1.6 mg/ml collagenase type I in HBSS without Ca^2+^ and Mg^2+^ (Gibco) at 37 %C for 25-30min.

### Kidney immunohistochemistry

One side kidney was taken after mice were killed by Cervical detachment at the 5th, 10th, 15th, 20th, 25th, and 30th day. The kidney was fixed for 2-3 days in 4% paraformaldehyde for paraffin sectioning. The slice thickness is about 4 microns. Sections were stained with hematoxylin-eosin and then observed under a microscope. The area of fibrosis and the number of inflammatory cells were counted by HC Image software.

### HEK-293T culture and transfection

We obtained HEK-293T cells from Dr. Lan Lei’s laboratory at Nanjing Normal University. The HEK-293T cells were cultured in growth medium that contained 90% DMEM, 10% FBS, 1% penicillin-streptomycin, at 37 °C in a humidified incubator with 5% CO2. The plasmids were transfected into the HEK-293T cells with Lipofectamine 2000 (Invitrogen) according to the manufacturer’s instructions.

### Calcium imaging

We performed calcium imaging experiments at room temperature (24-26 °C). Cells were loaded by Fura 2-AM (2μM, 15min, room temperature) before calcium imaging. HCimage software (HCimage Live 4.2 (Hamamatsu,Japan)) was used to control the hardware (TILL Polychrome V, German) and collected the data. The calcium imaging buffer consisted of (in mM): NaCl, KCl, CaCl_2_•2H_2_O, Glucose, HEPES, MgCl_2_•6H_2_O, NaH_2_PO_4_•2H_2_O.

### Whole-cell patch clamp recording

Whole-cell current clamp recordings were performed in DRG neurons. The current measurements were performed with an Axon 700B amplifier and Pclamp10.2 software package (Axon Instruments®). The electrodes were pulled from borosilicate glass (WPI Inc.). The electrode resistances was 2-4MΩ. All experiments were performed at room temperature (~26 °C). The DRG neurons were perfused with extracellular solution consisting of (in mM): NaCl 140, KCl 4, CaCl_2_ 2, MgCl_2_ 2, HEPES 10, glucose 5, with pH adjusted to 7.38 using NaOH. The intracellular pipette solution contained (in mM): KCl 135, MgATP 3, Na_2_ATP 0.5, CaCl_2_ 1.1, EGTA 2, glucose 5, with pH adjusted to 7.38 using KOH and osmolarity adjusted to 300 mOsm using sucrose.

### Samples preparation

Human serum was obtained from the Jiangsu Provincial Hospital of Traditional Chinese Medicine. Serum samples were thawed at 4 °C before experiments as in earlierresearch [43, 54]. Acetonitrile (HPLC grade, TEDIA, USA) was added into each serum sample (v/v=3:1) to precipitate protein. After violent shaking for 30s, the mixed samples were stewed at room temperature for 10min. Then, the mixture was centrifuged at 12000rpm two times for 10 min. Supernatant was prepared for UPLC-QTOF/MS. The kidney, urine and serum of the mice were treated in the same way. Finally, about 4μL supernatant was used for UPLC-QTOF/MS.

### UPLC-QTOF/MS

UHPLC analysis was performed using a ACQUITY UHPLC system (Waters Corporation, Milford, USA). Chromatographic separation was performed on a Thermo Syncronis C18 column (2.1 mm i.d. × 100 mm, 1.7 μm). The mobile phase contained solvent A (waters consist of 5mM ammonium formate, ammonium acetate, 0.2% formic acid) and solution B (acetonitrile contained 1mM ammonium formate, ammonium acetate, 0.2% formic acid) under gradient elution conditions: 10% A for 0-3min, 10-18% A for 3-9min, 18-20% A for 9-15min, 20-46% A for 16-18min, 46% A for 18-20min. The flow rate was controlled at 0.4 mL/min, and the column temperature was maintained at 30 %C. The mass spectrometry conditions were as follows: capillary voltage 3.0kV, source temperature 120 °C, desolvation temperature 350 %C, cone voltage 45V.

### Statistical analysis

All of the results are presented as means ± SEM. The data were statistically analyzed with two-tailed, paired/unpaired Student’s t test and a one-way or two-way ANOVA. Significance was considered as *P*<0.05

## Acknowledgments

This work was supported by the National Science Foundation of China to Z-X.T (31471007, 31771163)

